# The comparable strategic flexibility of model-free and model-based learning

**DOI:** 10.1101/2019.12.28.879965

**Authors:** Alexandre L. S. Filipowicz, Jonathan Levine, Eugenio Piasini, Gaia Tavoni, Joseph W. Kable, Joshua I. Gold

## Abstract

Different learning strategies are thought to fall along a continuum that ranges from simple, inflexible, and fast “model-free” strategies, to more complex, flexible, and deliberative “model-based strategies”. Here we show that, contrary to this proposal, strategies at both ends of this continuum can be equally flexible, effective, and time-intensive. We analyzed behavior of adult human subjects performing a canonical learning task used to distinguish between model-free and model-based strategies. Subjects using either strategy showed similarly high information complexity, a measure of strategic flexibility, and comparable accuracy and response times. This similarity was apparent despite the generally higher computational complexity of model-based algorithms and fundamental differences in how each strategy learned: model-free learning was driven primarily by observed past responses, whereas model-based learning was driven primarily by inferences about latent task features. Thus, model-free and model-based learning differ in the information they use to learn but can support comparably flexible behavior.

**Statement of Relevance:** The distinction between model-free and model-based learning is an influential framework that has been used extensively to understand individual- and task-dependent differences in learning by both healthy and clinical populations. A common interpretation of this distinction that model-based strategies are more complex and therefore more flexible than model-free strategies. However, this interpretation conflates computational complexity, which relates to processing resources and generally higher for model-based algorithms, with information complexity, which reflects flexibility but has rarely been measured. Here we use a metric of information complexity to demonstrate that, contrary to this interpretation, model-free and model-based strategies can be equally flexible, effective, and time-intensive and are better distinguished by the nature of the information from which they learn. Our results counter common interpretations of model-free versus model-based learning and demonstrate the general usefulness of information complexity for assessing different forms of strategic flexibility.

## Introduction

Humans can adopt a broad range of flexible strategies that learn from past observations to make accurate predictions about the future (Filipowicz, Anderson, & Danckert, 2016; Nassar, Wilson, Heasly, & Gold, 2010; O’Reilly, 2013; Stöttinger, Filipowicz, Danckert, & Anderson, 2014; Tenenbaum, Kemp, Griffiths, & Goodman, 2011). A prominent framework for capturing some of this range of strategies in humans is based on a distinction between model-free and model-based reinforcement learning (Daw, Niv, & Dayan, 2005; Decker, Otto, Daw, & Hartley, 2016; Eppinger, Walter, Heekeren, & Li, 2013; Gläscher, Daw, Dayan, & O’Doherty, 2010; Kool, Cushman, & Gershman, 2016; Pauli, Cockburn, Pool, Pérez, & O’Doherty, 2018; Sutton & Barto, 1998). Model-free strategies learn action policies that aim to maximize reward using only past stimulus-action outcomes. In contrast, model-based strategies learn action policies using both past stimulus-action outcomes and explicit representations of learned, latent statistical properties of the environment (e.g., transition probabilities between different states). Tendencies towards either strategy have been used as measures of individual learning traits. Tendencies towards model-free strategies have been equated with propensities towards low-level habit formation and are often interpreted as relatively automatic, habitual and strategically inflexible. Conversely, tendencies towards model-based strategies have been equated with more deliberate, goal-directed, and flexible learning abilities (Daw et al., 2005; Gillan, Otto, Phelps, & Daw, 2015; Gläscher et al., 2010; Kool, Gershman, & Cushman, 2017, 2018; Pauli et al., 2018). In general, people tend to use a mix of model-free and model-based strategies, which can depend on the specific task conditions (Kim, Park, O’Doherty, & Lee, 2018; Kool et al., 2017) and individual differences in age (Decker et al., 2016; Eppinger et al., 2013) and psychiatric symptoms (Sebold et al., 2014; Voon et al., 2015).

The idea that model-based strategies are more flexible than model-free strategies seems to have arisen from the observation that, in general, model-based strategies are more computationally complex than model-free strategies. Computational complexity corresponds to the computational resources (e.g., computational time, memory) required to implement an algorithm, and how these resources scale with increased input. Resource demands depend on the algorithm used to implement a specific learning strategy and can be computed in different ways (Bossaerts & Murawski, 2017; Bossaerts, Yadav, & Murawski, 2019; Kool et al., 2018; Polonio, Di Guida, & Coricelli, 2015). These resource demands are generally lower for model-free algorithms, which track only stimulus-action outcome mappings, than for model-based algorithms, which must also learn, update, and use representations of other task-relevant variables. As such, this language has begun to permeate the human model-based and model-free decision-making literature, such that model-based strategies are often described as having more strategic ‘sophistication’ or ‘complexity’ than model-free strategies, which is equated with more flexible behavior (Decker et al., 2016; Doll, Simon, & Daw, 2012; Kim et al., 2018; Kool et al., 2018).

However, computational complexity is not an appropriate measure of strategic flexibility. Strategic flexibility concerns the relationship between inputs (observations from the environment) and outputs (behavioral responses). In general, the more forms that this relationship can take, the more flexible the behavior. Computational complexity does not explicitly measure this type of flexibility. For example, the exact same strategy, with the same relationship between input and output, can be implemented by different algorithms that use different computational resources more or less efficiently (which is a focus of the field of optimization in computer science; Cormen, Leiserson, Rivest, & Stein, 2009). Instead, strategic flexibility is better measured by statistical or information complexity (Bialek, Nemenman, & Tishby, 2001; Grünwald & Rissanen, 2007; Myung, Balasubramanian, & Pitt, 2000). Unlike computational complexity, statistical complexity depends directly on the relationship between input and output, reflecting how flexibly a model or algorithm changes its behavior (output) when trained on different patterns of data (input). This form of complexity can be estimated in decision-making tasks using a method inspired by the information-bottleneck, which measures the amount of past information subjects use to make decisions (Tishby, Pereira, & Bialek, 2000). In general, strategies that incorporate more past information are regarded as being more flexible, because this increased information indicates a greater diversity in how past observations influence current behavior (Filipowicz, Glaze, Kable, & Gold, 2020; Gilad-Bachrach, Navot, & Tishby, 2003; Glaze, Filipowicz, Kable, Balasubramanian, & Gold, 2018). Thus, while model-based learning strategies may be more computationally complex, this does not guarantee that they are more information complex, and therefore does not also guarantee that they are more flexible.

The goal of this study was to measure the strategic flexibility of model-based versus model-free learning in terms of their information complexity. To achieve this goal, we estimated information complexity from behavioral choice data of human subjects performing a canonical task used to distinguish these strategies (Daw, Gershman, Seymour, Dayan, & Dolan, 2011). We used an information-based measure that was applied directly to each subject’s choice data and did not require explicit knowledge of the strategy they used to generate those choices. Contrary to common descriptions of model-based strategies as more complex and more flexible, we show that both model-based and model-free strategies can show similar degrees of information complexity on this task, with comparable levels of accuracy and response times (RTs). We further show that this information-based approach identifies a more fundamental distinction between these types of strategies, which is the nature of the feature spaces over which they perform inference: observable features for model-free learning and latent features for model-based learning.

## Methods

### Participants and behavioral task

We assessed information complexity from behavioral data of human subjects performing the two-step task (Daw et al., 2011), which is commonly used to measure model-based and model-free learning. The data were obtained from a publicly available dataset consisting of 197 subjects, recruited on Amazon Mechanical Turk, who performed the two-step task that Kool and colleagues describe as the “Daw two-step task” (see Kool et al., 2016 for full task and subject details; data available at https://github.com/wkool/tradeoffs). For this task, subjects choose one of two first-step actions that each lead stochastically to one of two second-step states. Each first-step action leads distinctly to one of the two possible second-step states with 0.7 probability (common transition) and to the other state with 0.3 probability (rare transition; Fig. 1a). Subjects then choose between two additional actions in this second state, which lead stochastically to a reward. Reward probabilities for each second-step action drift independently according to a Gaussian random walk (μ=0, σ=0.025) with reflecting boundaries at 0.25 and 0.75 (Fig. 1b). A subject’s goal in this task is to accumulate as much reward as possible.

**Figure 1.**
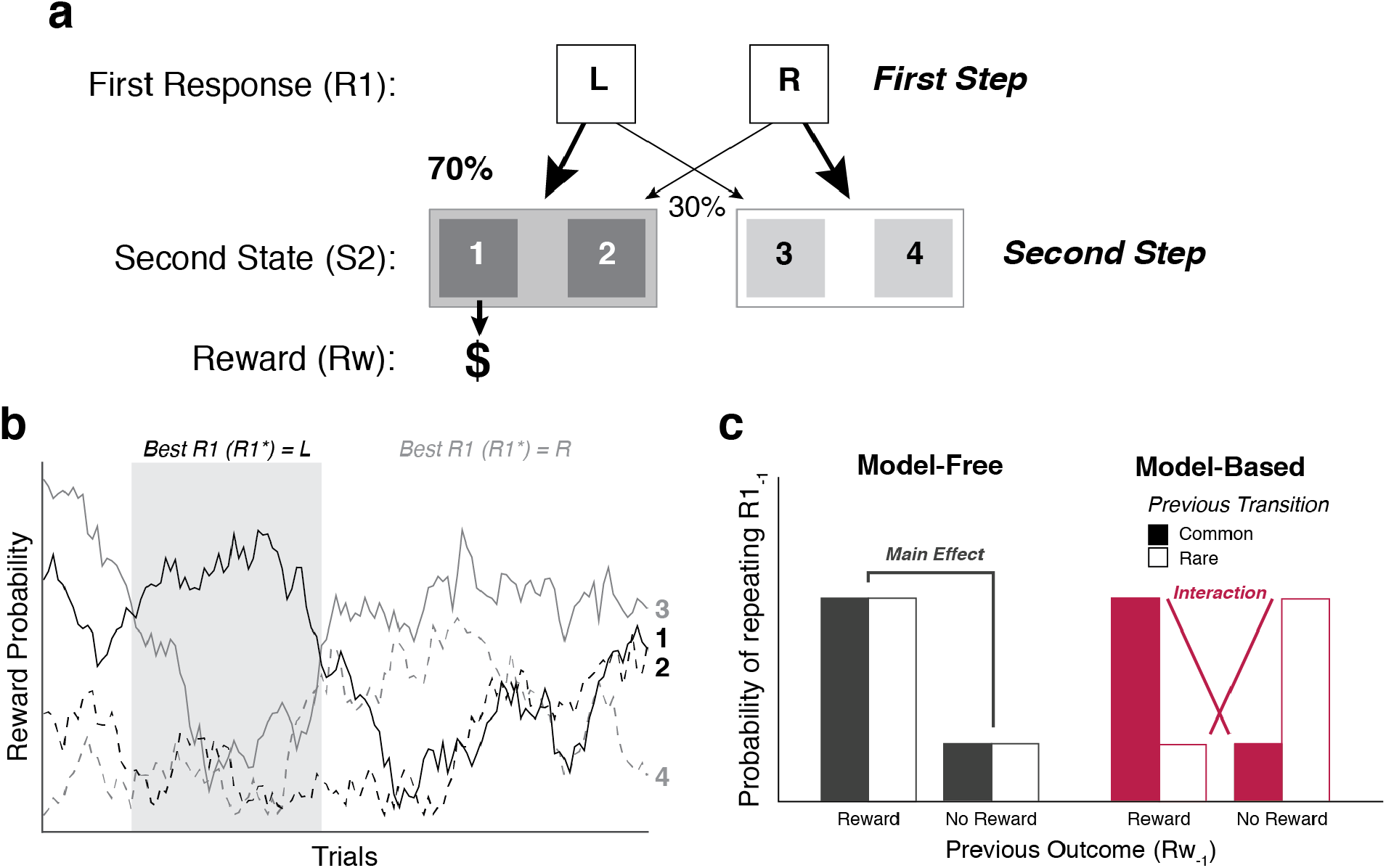
Description of the two-step task. (a) The subject makes a first-step choice between two alternatives (L or R) that lead probabilistically to one of two second-step states (filled or open rectangles). Subjects then make a second choice that leads probabilistically to a reward. (b) Example reward probabilities for each second-step option, which drift independently according to a Gaussian random walk. (c) Examples of response patterns for model-free and model-based strategies. Model-free strategies tend to repeat first-step choices that lead to a reward, ignoring the transition structure of the environment. A behavioral signature of these strategies is a main effect of reward on the probability of repeating the same first-step choice as the previous trial. Model-based strategies use the transition structure of the environment to make choices. A behavioral signature of model-based strategies is an interaction between reward and previous transition type (common or rare) on the probability of repeating the same first-step choice as the previous trial.

This task was designed to identify propensities towards more model-based or model-free strategies (Daw et al., 2011; Kool et al., 2016). Purely model-free strategies rely solely on the observed stimulus-action mappings without attempting to learn or use latent information about the transition between the first and second steps. As a result, model-free strategies tend to repeat first-step actions that lead to second-step reward, regardless of whether the first to second-step transition was rare or common. This tendency can be measured behaviorally as a main effect of reward on the probability of repeating a previous first-step action (Fig. 1c). In contrast, model-based strategies account for the latent transition structure of the task environment and therefore select first-step actions that maximize the chance of returning to a rewarding second-step state. This tendency can be measured behaviorally by the interaction of reward and transition type (rare or common) on the probability of repeating a first-step action (Fig. 1c).

All available data were used for our analyses without any additional exclusions. As outlined in the original article, the original study was approved by the Harvard Committee on the Use of Human Subjects and all subjects gave informed consent.

### Measures of information complexity and predictive accuracy

We computed the information complexity and predictive accuracy of each subject using a method inspired by the information-bottleneck (Tishby et al., 2000). This method assumes that subjects form an internal belief or model *M* from past task observations (*X*_*past*_) to predict some future aspect(s) of the task (*X*_*future*_). The amount of information *M* encodes from *X*_*past*_ is measured by their mutual information; i.e., *I*_*past*_ = *I*(*X*_*past*_; *M*). Larger values of *I*_*past*_ correspond to models with higher information complexity (Filipowicz et al., 2020; Gilad-Bachrach et al., 2003; Glaze et al., 2018). Predictive accuracy was measured as the mutual information between the model and future observations; i.e., *I*_*future*_ = *I*(*M*; *X*_*future*_). Larger values of *I*_*future*_ correspond to models with high predictive accuracy (Gilad-Bachrach et al., 2003; Palmer, Marre, Berry, & Bialek, 2015; Tishby et al., 2000). The main strength of this information-based measure is that it quantifies information complexity without requiring any explicit knowledge of the strategy producing the behavior.

To compute *I*_*past*_, we first identified the four observed and latent variables that in principle could be used to perform the task: 1) the first-step response (*R*1; a directly observed quantity), 2) the second-step transition (*S*2; a directly observed quantity), 3) the reward after second-step response (*Rw*; a directly observed quantity), and 4) the firsts-tep response that would maximize the chance of obtaining a reward (*R*1^*^; a latent quantity). These four variables were combined for each trial to form past trial “features” (*F*), which could take 16 unique possible values. Information complexity was computed for each subject by measuring the mutual information between the features from the previous trial *F*_−1_ and the first-step responses on the current trial *R*1_0_:

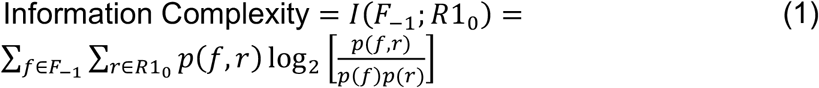

Predictive accuracy was computed for each subject as the mutual information between subject first-step responses (*R*1_0_) and the aspect of the task they were attempting to predict, 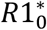, which corresponds to the best action to take, given the current task contingencies, to maximize reward (Fig. 2a):

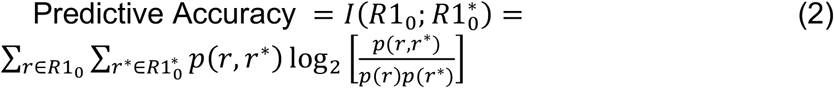

**Figure 2.**
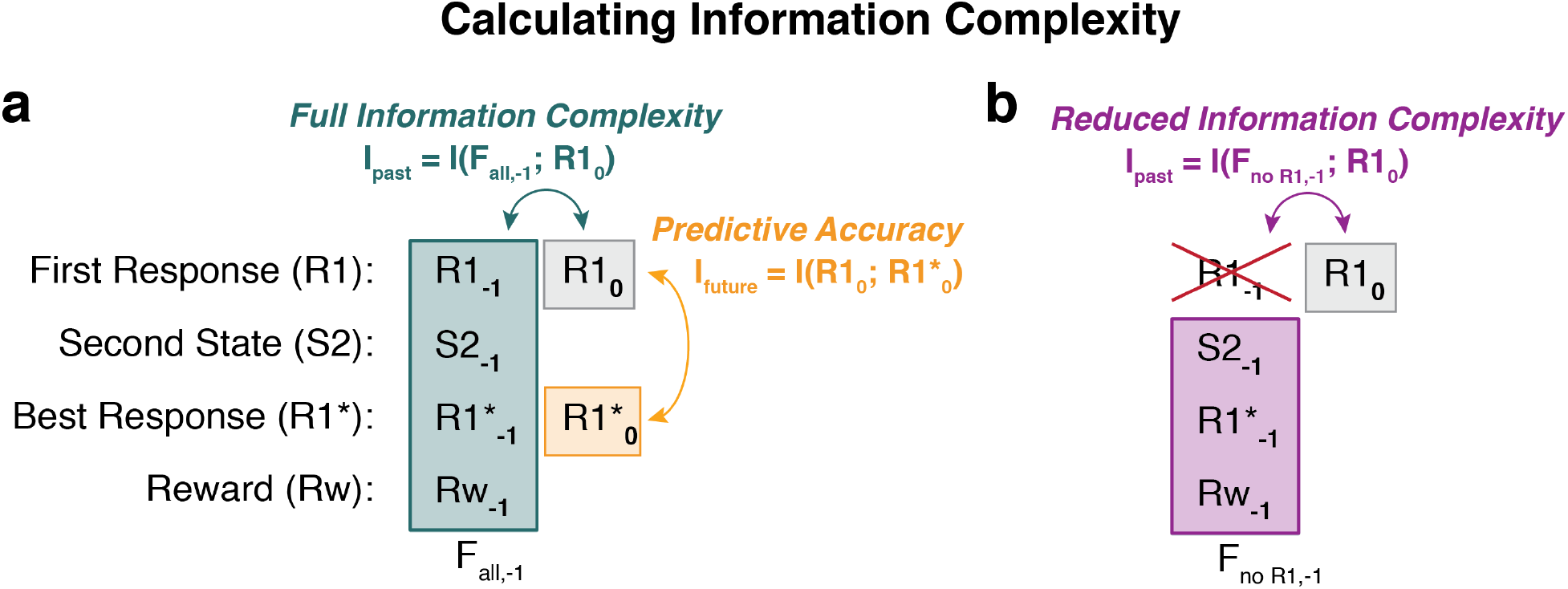
Calculation of information complexity. (a) Full information complexity was measured as the mutual information between task features one trial in the past (*F*_*-1*_, including first-step responses, *R1*_*-1*_, second-step transitions, *S2*_*-1*_, reward received, *Rw*_*-1*_, and best first responses, 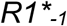) and current responses (*R1*_*0*_). Predictive accuracy was measured as the mutual information between *R1*_*0*_ and 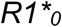. (b) Reduced information complexity was measured as the mutual information between all except one element from the previous trial and subject responses (*R*1_0_). This example shows the calculation of reduced complexity when omitting past first-step responses.

This measure of predictive accuracy captures the effectiveness of first-step responses; i.e., how well subjects are making responses that will most likely lead them to the best second-step states.

Given the Markovian nature of both the process generating the stimuli and the processes guiding the models that are generally used to capture subject strategies in this task, past features included only elements of the previous trial and did not extend further in the past. Similar to previous applications of this method to experimental data, we also assumed that subjects treated the task in a Markovian manner, by including the latent variable 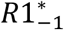 in the past-feature vector as a proxy for the history of previously observed transitions (Filipowicz et al., 2020; Glaze et al., 2018). We also omitted the second-step response from the past-feature vector, because the information provided by this past element does not inform the first-step responses in simulations beyond the information provided by the combination of the second-step transition (***S2***) and the reward (***Rw***), and including this element in the past-feature vector did not improve our ability to distinguish between simulations of model-based and model-free agents. Omitting the second-step response also helps estimate mutual information more accurately, by reducing the size of the joint distribution between the past features and first-step responses (*p*(*F*_−1_, *R*1_0_) and thus limiting misestimates due to the “curse of dimensionality” (Bellman, 1961).

Complexity and accuracy can also be computed with respect to second-step responses (*R*2) and best second-step choices (*R*2^*^). However, the response policies governing these choices are generally assumed to be identical across strategies. Moreover, previous analyses have primarily concentrated on first-step responses, because these responses provide the clearest distinction between model-free and model-based strategies. Therefore, because of the dimensionality issues highlighted above, we chose to omit these features from our analyses.

### Behavioral metrics of model-based and model-free learning

A common behavioral metric of model-based and model-free learning is to measure each subject’s main effect of reward and reward-by-transition-type interaction (Fig. 1c). Each subject’s main effect of reward was computed as the proportion of trials on which the subject repeated the same first-step response as the previous trial after receiving a reward minus the proportion of repeated first-step responses when no reward was received (independent of second-step transition). Each subject’s reward-by-transition-type interaction was computed as the proportion of trials on which the subject repeated first-step responses after reward/no reward on previous common/rare transitions, respectively, minus the proportion of repeated first-step responses after no reward/reward on previous common/rare transitions, respectively (Kool et al., 2016).

### Computational models

We fit the model-based and model-free learning algorithms used in (Kool et al., 2016) to each subject’s choice data. The model-free algorithm is based on a SARSA(λ) temporal-difference learning algorithm that updates Q-values (*Q*_*MF*_) of stimulus-action pairs (*s*_*i,j*_, *a*_*i,j*_), where *i* indicates the step (1 or 2) and *j* denotes the state (used only for second-step states after *a*_1_ in *s*_1_ is taken, given that there is only one first-step state). The Q-values at the first-step are updated according to:

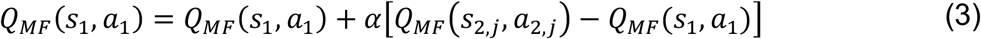

where *α* corresponds to a fixed learning rate that determines the extent to which current values are modified by the reward prediction error, which at the first step is driven by the difference between the value of the action taken at the second step and the action taken at the first step *Q*_*MF*_(*s*_2,*j*_, *a*_2*j*_) − *Q*_*MF*_(*s*_1_, *a*_1_). When outcomes are observed at the second step, the Q-value for the second-step action is updated using the observed reward (*r*_2_):

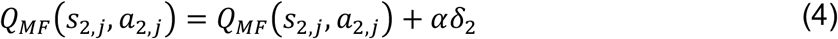

where

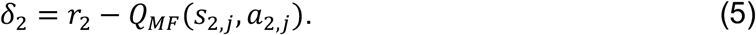

The first level Q-values are then updated again as a function of the second-step prediction error weighted by an eligibility trace decay parameter *λ* such that when *λ* = 0, only the values of second steps get updated:

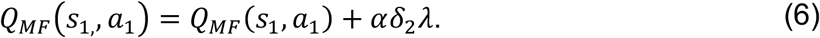

The model-based algorithm uses the transition probability structure of the task to select actions. First-step model-based Q-values (*Q*_*MB*_) are calculated using Bellman’s equation, which uses the expected values of second-step actions and the transition structure of the environment to determine expected values for each of the first-step actions:

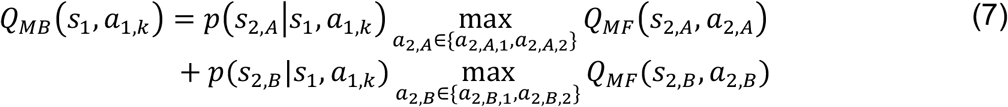

which is performed for the *k* possible first-step actions. Q-values at the second step are updated in the same manner as the model-free algorithm. Therefore, these strategies differ only in the computations they use to determine their first-step action values.

We simulated two different model-based agents to measure the influence of different past task features on information complexity. The first, “stochastic” model-based agent knew and used the actual transition probabilities between first-step actions and each second-step state (0.70 for common and 0.30 for rare). The second, “deterministic” model-based agent treated the transitions between first-step actions and second-step states as deterministic, believing that each first-level action always transitioned to distinct second-step states.

To measure the degree to which subjects mixed model-based and model-free action policies, a strategy-mixing coefficient (*w*) was added to the model, the value of which varies from 0 (a fully model-free strategy) to 1 (a fully model-based strategy):

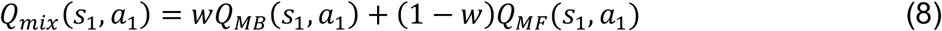

For all models, Q-values at each step were converted into action probabilities by applying a softmax function:

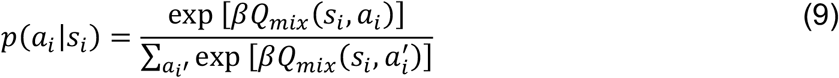

where *β* corresponds to an inverse temperature parameter that controls the randomness of the choice as a function of the Q-values (i.e., as *β* → 0 action probabilities tends to become uniform, and as *β* → ∞ the probability of choosing the action with the highest Q-value tends towards 1).

Similar to Kool and colleagues (2016), the models we fit to subject data were identical to the mixture model outline above, with two parameters added to the first-step softmax decision rule:

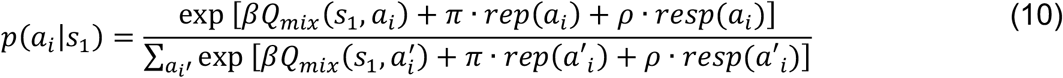

where *π* indicates the degree of perseveration, or ‘stickiness’, towards repeating the same motor action as on the previous trial (with *π* > 0 indicating perseveration), and *ρ* indicates perseveration towards the same first-step option (i.e., the fractal that was selected). Perseveration for first-step actions and choices could be distinguished because the action required to select each option was randomly chosen on each trial. Models were fit using the same model fitting procedures outlined by Kool and colleagues (2016). Briefly, we used the *mfit* toolbox (Gershman, 2016) to obtain maximum *a posterior* parameter estimates obtained by gradient decent using the same weakly informative priors used by Kool and colleagues.

### Model simulations

For our simulations, we performed 100 simulations of 10,000 trials for values of *w* (from eq. 8) that varied between 0 and 1 in increments of 0.1. Information metrics for each value of *w* were averaged across all 100 simulations. Apart from differences in the *w* mixing parameter, all other parameters were kept constant in each simulation, consistent with the values used by Kool and colleagues (*α* = 1, *β* = 5, *λ* = 0.5). Although the specific information values fluctuated with different parameter values, all of the trends reported in the manuscript held for a wide range of *α* and *β* values, except when *α* → 0 (no learning occurs) or *β* → 0 (behavior is random).

## Results

### Model-free and model-based strategies are similarly flexible and effective

As was reported previously, individual subjects performing the two-step-task differed considerably in their propensity to use more model-free or model-based strategies (Kool et al., 2016). To quantify this propensity, we computed each subject’s main effect of reward on the probability of repeating the same first-step action as the previous trial (a metric of model-free tendencies), and the interaction between reward and transition type (rare or common; a metric of model-based tendencies; Fig. 1c). We then compared these data-driven metrics of model-free and model-based tendencies to each subject’s strategic complexity using our method inspired by the information-bottleneck (Filipowicz et al., 2020; Gilad-Bachrach et al., 2003; Glaze et al., 2018; Tishby et al., 2000; Fig 2a), which also is data-driven and thus did not require any explicit assumptions about the specific strategy each subject used to perform the task. Because the main effect and interaction terms were negatively correlated with each other (Spearman’s rho=-0.26, *p*=0.0002), we computed semi-partial Spearman correlations between complexity and each variable (main effect and interaction), while accounting for the other variable.

Subjects with either higher main effects (i.e., greater model-free tendencies) or higher interactions (i.e., greater model-based tendencies) tended to have higher information complexity (Fig. 3). Therefore, subjects with choices that were more consistent with either model-based or model-free strategies had relatively high information complexity.

**Figure 3.**
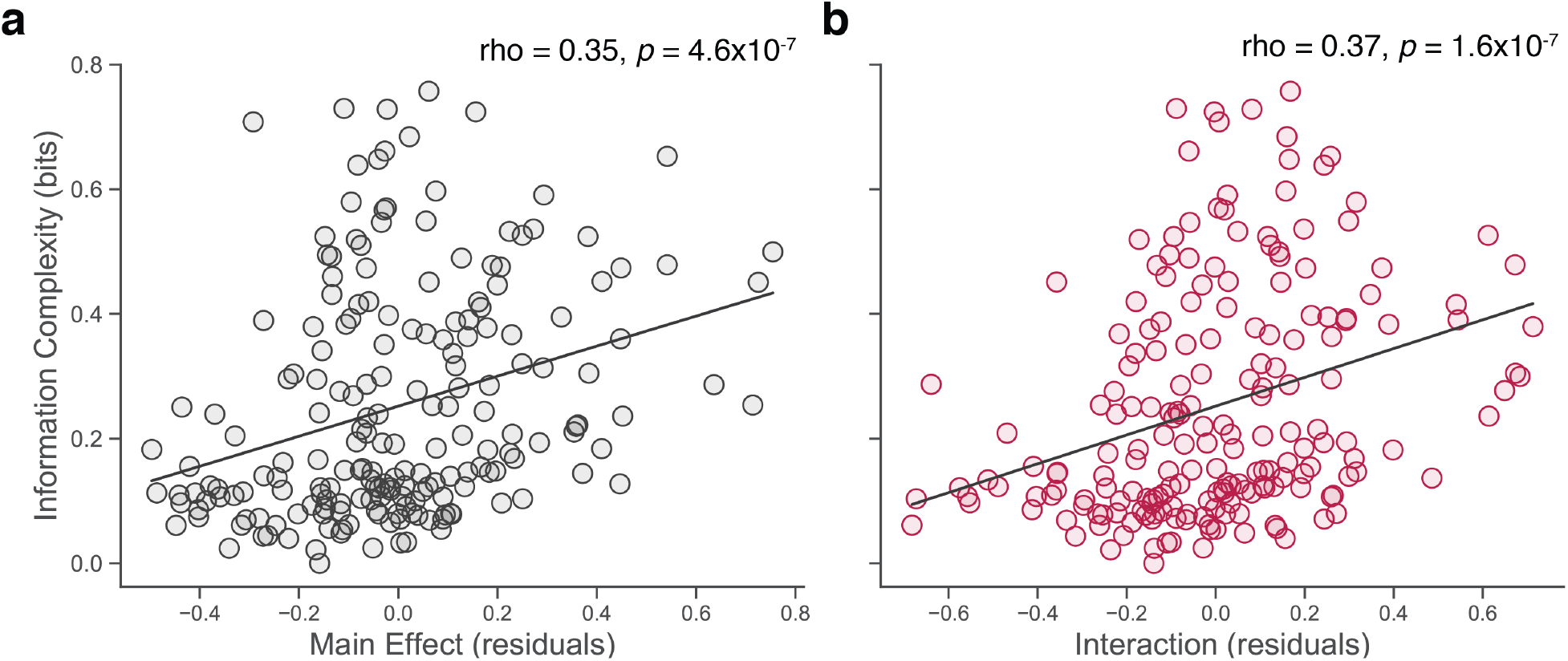
Information complexity was correlated positively with the size of both (a) main effect (a measure of the tendency to use a model-free strategy) and (b) interaction (a measure of the tendency to use a model-based strategy) values (see Fig. 1c) from individual subjects (points). *rho* values are semi-partial Spearman correlation coefficients of per-subject information complexity versus main effect (a) or interaction (b) values while accounting for the other value; *p*-values are for *H*_0_: *rho*=0. Plotted points represent residuals after accounting for interaction (a) and main effect (b) values. Lines are least-squares linear fits.

Consistent with the trends observed by Kool and colleagues (2016), increases in information complexity did not necessarily improve subject outcomes on the task: subjects using more information complex strategies did not show reliable increases in obtained reward (Fig. 4a). Likewise, increased tendencies towards model-free or model-based learning strategies, as measured by main effect and interaction values respectively, were not associated with increases in reward (Fig. 4d,g).

**Figure 4.**
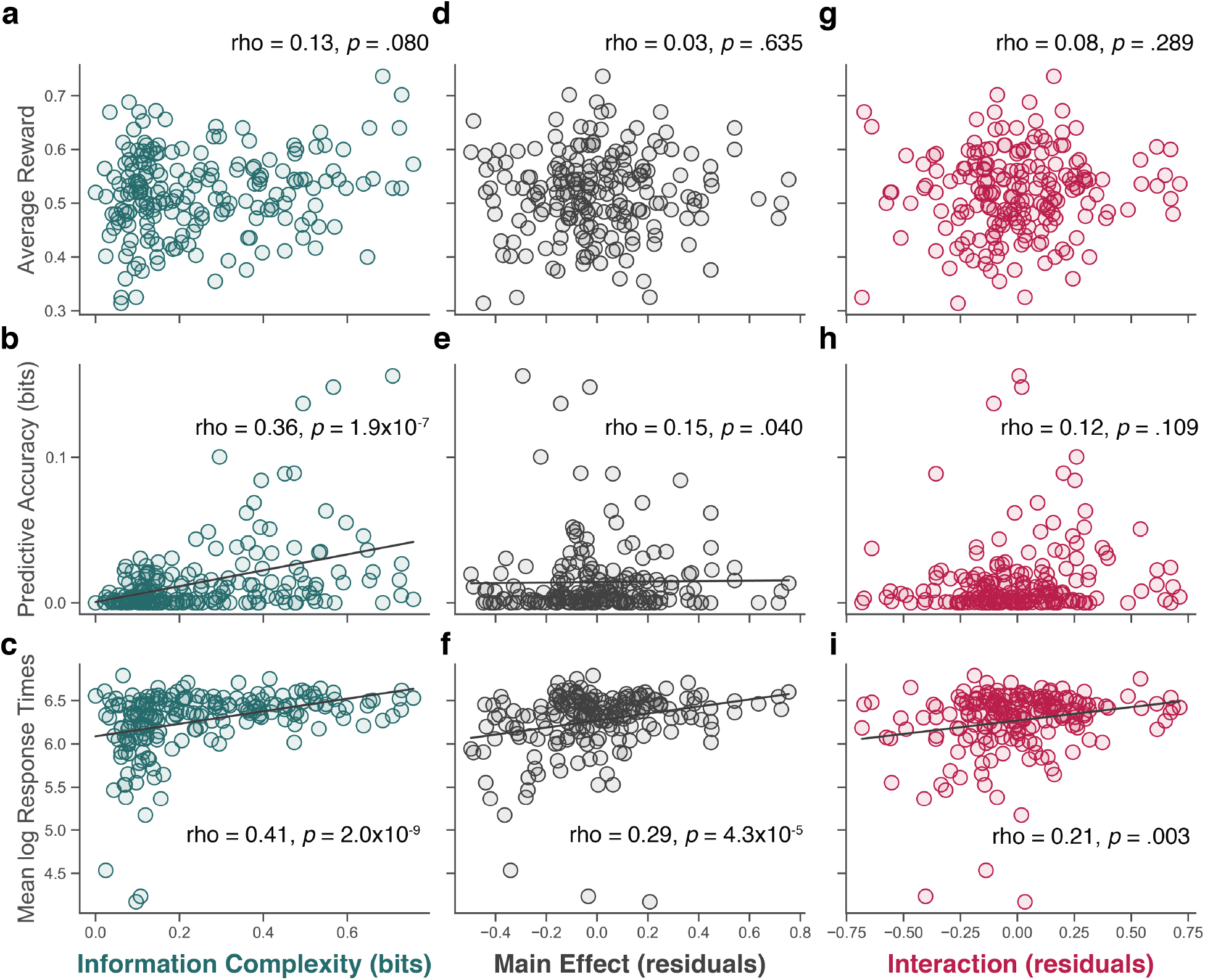
Increasing information complexity for individual subjects (points) did not correspond to increases in average reward (a) but did correspond to increases in predictive accuracy (b) and longer RTs (c). Increased model-free behavior (i.e., higher main effects) corresponded with similar tendencies (d-f), and increased model-based behavior (i.e., higher interactions) corresponded to increases in RTs, but not average reward or predictive accuracy (g-i). Correlation values for main effect and interaction values indicate semi-partial Spearman correlation coefficients while accounting for the other value, respectively. Plotted main effect and interaction values in (d–i) correspond to residuals after accounting for interaction and main effect values, respectively. Lines are linear least-squares fits and plotted for Spearman correlations with *p*<.05.

In contrast, subjects with higher information complexity tended to have higher predictive accuracy, which implies that more-complex subjects made responses that were better tuned to the first-step response that would maximize their chances of obtaining a reward (Fig. 4b). This discrepancy with average reward can be explained by the relatively low information shared between the most rewarding first-step action and the actual probability of being rewarded for taking this action. For example, consider an extreme case in which one second-step state includes an option that offers the maximum possible 0.75 reward probability. In this case, a subject that consistently makes the best first-step response will have a reward probability of only 0.53 (i.e., the transition probability of 0.7 times the reward probability of 0.75). Thus, for this task using strategies higher in information complexity did not provide any advantages in terms of reward obtained but did improve predictive accuracy. This relationship was not driven by a particular tendency to use a model-free or model-based strategy, neither of which was related strongly with predictive accuracy (Fig. 4e,h).

There were, however, similar across-subject relationships between response times (RTs) and either information complexity, a tendency to use a model-free strategy (main effect), or a tendency to use a model-based-strategy (interaction). Overall, mean log RT, measured for each subject across all trials, tended to increase systematically as a function of the information complexity of a subject’s strategy (Fig. 4c), the main effect (Fig. 4f), and the interaction term (Fig. 4i). The relationship with model-free tendencies was particularly striking, given that model-free strategies are commonly framed as automatic, habitual type responses, which should predict shorter RTs for subjects using more model-free strategies. Instead, we found that increased use of either model-free or model-based strategies was associated with a systematic increase in RTs.

### Model-free and model-based strategies learn from different task features

Despite their strong similarities in information complexity, subjects using model-free versus model-based strategies learned from different combinations of features to solve the task. To examine the influence of specific feature elements on individual subject strategies, we computed information complexity between subject responses and vectors of past features that omitted each one of the four past elements we considered (Fig. 2a). We then subtracted this ‘reduced’ information complexity from each subject’s full information complexity to assess the extent to which their complexity was driven by each individual element (Fig. 5a).

**Figure 5.**
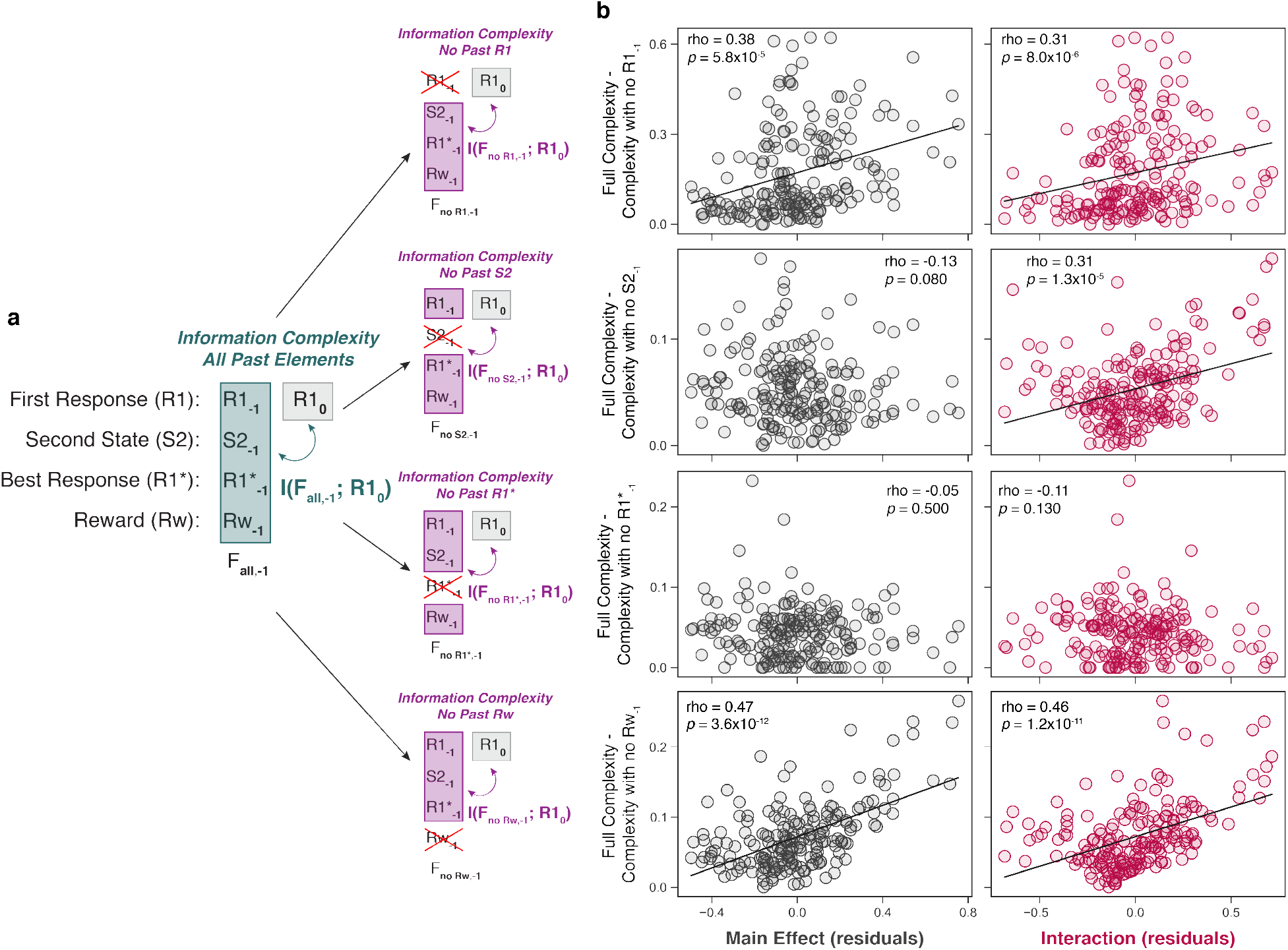
Contribution of different past features to subject information complexity. (a) The contribution of each individual past element (first response, *R1*; second-step state, *S2*; best response, *R1\**; reward, *Rw*) to overall information complexity was measured by subtracting the information complexity computed with each element removed from the past-feature vector (*F*_*−1*_) from the full information complexity computed using all of the past elements. (b) This difference in complexity was correlated with the individual subject (points) main effect (grey) and interaction (red) values, with higher values on the ordinate indicating stronger contribution of each element to subject complexity. Abscissa main effect and interaction values represent residuals for individual subjects after accounting for interaction and main effect values respectively. Points are data from individual subjects. Lines are least-squares linear fits and plotted for semi-partial Spearman correlations with *p*<.05.

The information provided by past first responses and past rewards was positively correlated with both main effect and interaction values, indicating that both of these elements contribute to the complexity of both model-free and model-based strategies (Fig. 5b). In contrast, the second-step state from the previous trial contributed more to the complexity of subjects with higher interaction values (i.e., that used more model-based strategies), but not main-effect values (i.e., that used more model-free strategies; Fig. 5b). Thus, subjects using more model-based or more model-free strategies were complex for different reasons: model-free strategies focused primarily on previous rewards and first-step responses, whereas model-based strategies used those elements in addition to information from the previous second-step state.

### Model-free and model-based algorithms show similar tendencies

To better understand how these complexity trends in the subjects’ behavior might relate to specific learning algorithms that have been proposed for model-based and model-free strategies, we measured the complexity of simulations using a commonly used computational model (Daw et al., 2011; Kool et al., 2016). This model assumes that agents use both model-based and model-free learning strategies that are mixed together with a coefficient *w*, where *w* = 0 corresponds to a purely model-free strategy, and *w* = 1 corresponds to a purely model-based strategy. We examined two versions of the model that differ in how the model-based strategy treats the transition probabilities in the task when calculating first-step action values: 1) a “stochastic-transition model”, in which the transition probability between first-step responses and second-step states is assumed to be known (equal to 0.7/0.3 in the task analyzed here, eq. 7); and 2) a “deterministic-transition model”, which acts as if the transitions between first-step actions and second-step states is deterministic, even if the true transition probabilities in the task are stochastic (see methods, eq. 7).

We first note that, consistent with the terminology used in computer science, the model-based algorithm is more computationally complex than the model-free algorithm. We defined computational complexity as the number of operations each algorithm uses to produce first-step actions values, because the purely model-based and model-free strategies differ only in the computation of these values. We chose a relatively conservative measure of computational complexity such that each use of addition, subtraction, multiplication, and argmax to derive a first-step action value counted as a single operation. Using this measure, a pure model-based strategy, which requires 10 operations to derive first-stage action values, is more computationally complex than a pure model-free strategy, which requires 6 operations. Using other approaches that take into account different computational costs of each operation (e.g., multiplication and argmax might require more computations for the brain to perform than operations such as addition or subtraction; Eliasmith & Anderson, 2003) would tend to further amplify this difference, because the model-based algorithm would incur more of such costs.

We next computed the information complexity for simulations that varied in the degree to which they were more model-based or model-free. Computing the information complexity for different values of *w*, we found that information complexity exhibited a non-monotonic decrease as the mixture moved away from either a pure model-free or pure model-based strategy, with minimum at *w*=0.8 for the stochastic transition model (Fig. 6a, green circles) and *w*=0.4 for the deterministic transition model (Fig. 6b, green circles). Information complexity was also higher for pure model-free than pure model-based strategies, though this difference was larger in the stochastic transition model (mean information complexity: model-free=0.05 bits, model-based=0.02 bits; Wilcoxon rank-sum test comparing information complexity between purely model-free and purely model-based simulations, *p*=2.5×10^-34^). The fact that these simulations, unlike the subject data, showed substantially lower information complexity for purely model-based strategies suggests that this computational model, and particularly the stochastic transition version which is typically used, does not capture the variety of model-based strategies that subjects actually use in the task, a discrepancy that has been noted previously (Akam, Costa, & Dayan, 2015; da Silva & Hare, 2020).

**Figure 6.**
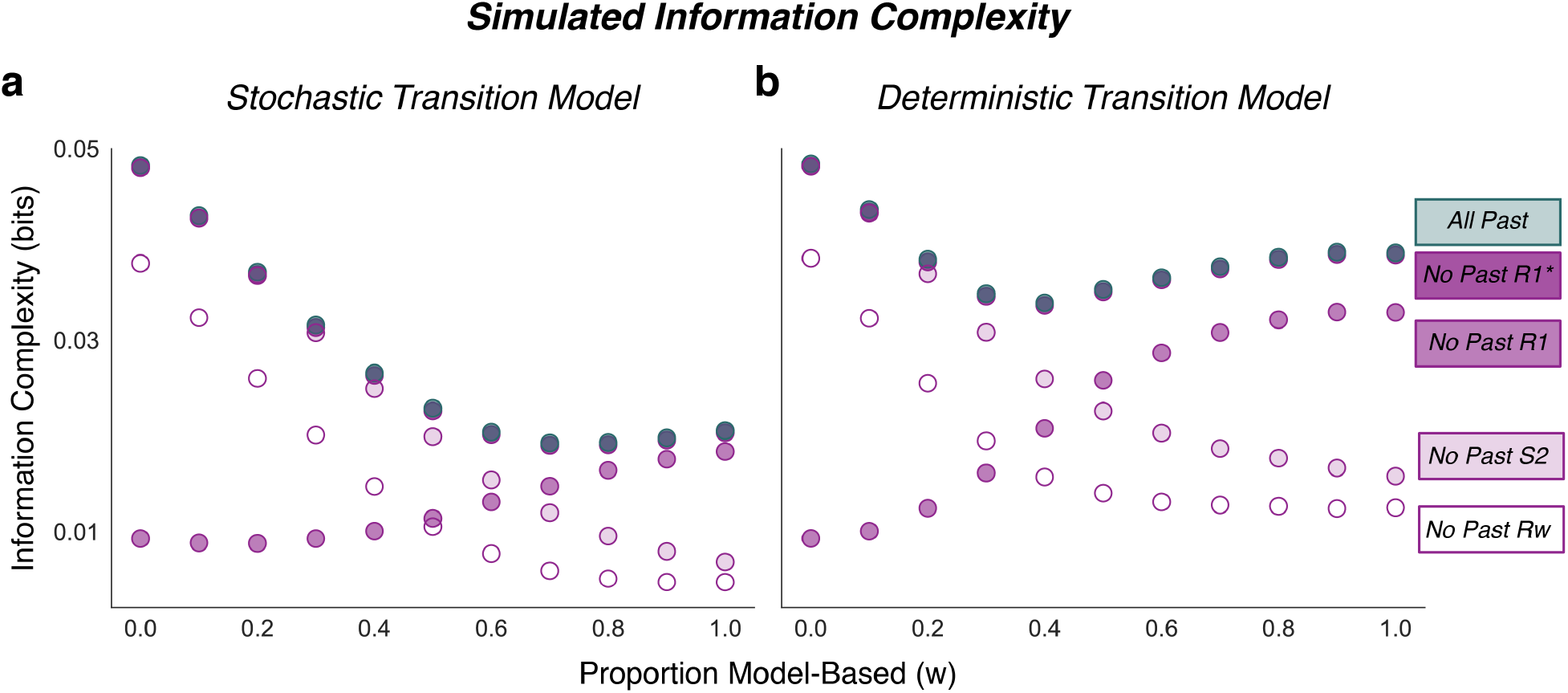
Information complexity for simulated model-free and model-based (either the stochastic-transition model in a or the deterministic-transition model in b) agents performing the two-step task. The simulations mixed model-free and model-based strategies using a strategy-mixing coefficient (*w*), with *w* = 1 indicating a fully model-based strategy. Information complexity was measured between all of the past features and simulated responses (green circle), along with reduced information complexity measured by omitting either the best first-step response, actual first-step response, second-step transition, or reward on the previous trial (purple circles, from darker to lighter fill, respectively; note that information values computed when omitting the previous best response are nearly entirely under the values of information complexity computed with all past features). Points indicate mean information values across 100 simulations for each value of *w*. SEM is smaller than size of the dots in all cases.

Simulations further reinforced the behavioral analyses from Fig. 5 showing that model-free and model-based strategies rely on different past features to perform the task. Like for the behavioral analyses, we computed the information complexity of simulated choices while removing each one of the four elements of the past-feature vector (Fig. 6). Omitting the previous first-step response (*R*1_−1_) resulted in a substantial decrease in information complexity that was particularly strong for more model-free agents (i.e., as *W* → 0; compare to the first row of Fig. 5b). This result reflects the fact that the model-free algorithm makes first-step choices according to action values that depend on whether the first action from the previous trial resulted in reward, regardless of whether this action led to a common or rare second-step transition. Conversely, omitting the second-step state (*S*2_−1_) from the past-feature vector resulted in substantially reduced information complexity just for model-based agents (i.e., as *W* → 1; compare to the second row of Fig. 5b). Omitting the previous best first-step response (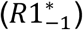) had little impact on model complexity regardless of the strategy (compare to the third row of Fig. 5b). Omitting the previous reward (*Rw*) reduced the complexity of both strategies (compare to the fourth row of Fig. 5b). These results indicate that the information complexity corresponding to a standard model-free algorithm is influenced heavily by the previous first response, whereas a standard model-based algorithm relies predominantly on information from the second-step transitions from the previous trial. The fact that subjects exhibiting model-based tendencies showed a strong influence of the previous first response on information complexity (Fig. 5b), whereas this influence was not seen as strongly in model-based simulations, further supports the idea that people use a diversity of model-based strategies not necessarily captured by these standard models (Akam et al., 2015; da Silva & Hare, 2020).

Consistent with these simulations and further refuting claims that strategic flexibility increases along a continuum from model-free to model-based learning, fits of these models to behavior did not show systematic, monotonic relationships with information complexity. Specifically, we fit to each subject’s behavioral choices a learning model that mixed both the model-free algorithm and the model-based algorithm (either the stochastic-transition model or the deterministic-transition model, which produced roughly equivalent fits: median BIC [interquartile range] across subjects for the stochastic-transition model=292 [241–337], and for the deterministic-transition model=292 [240– 337]; Wilcoxon signed-rank test for *H*_*0*_: median BIC difference=0, *p*=0.687), using the same strategy-mixing coefficient (*w*) as in the simulations above. Using the fits from these models, we found no correlation between the best-fitting value of *w* and information complexity in either the fits provided by the stochastic transition model (Fig. 7a) or the deterministic transition model (Fig. 7c). Additionally, we found no correlation between *w* from either model and predictive accuracy across subjects, suggesting that neither model-based nor model-free strategy type provided a distinct advantage for this task, which is consistent with previous reports that found no correlations between *w* and overall accumulated reward (Kool et al., 2016; Fig. 7a,c). We additionally found no correlation between the strategy-mixing coefficient from either model and RT (Fig. 7b,d).

**Figure 7.**
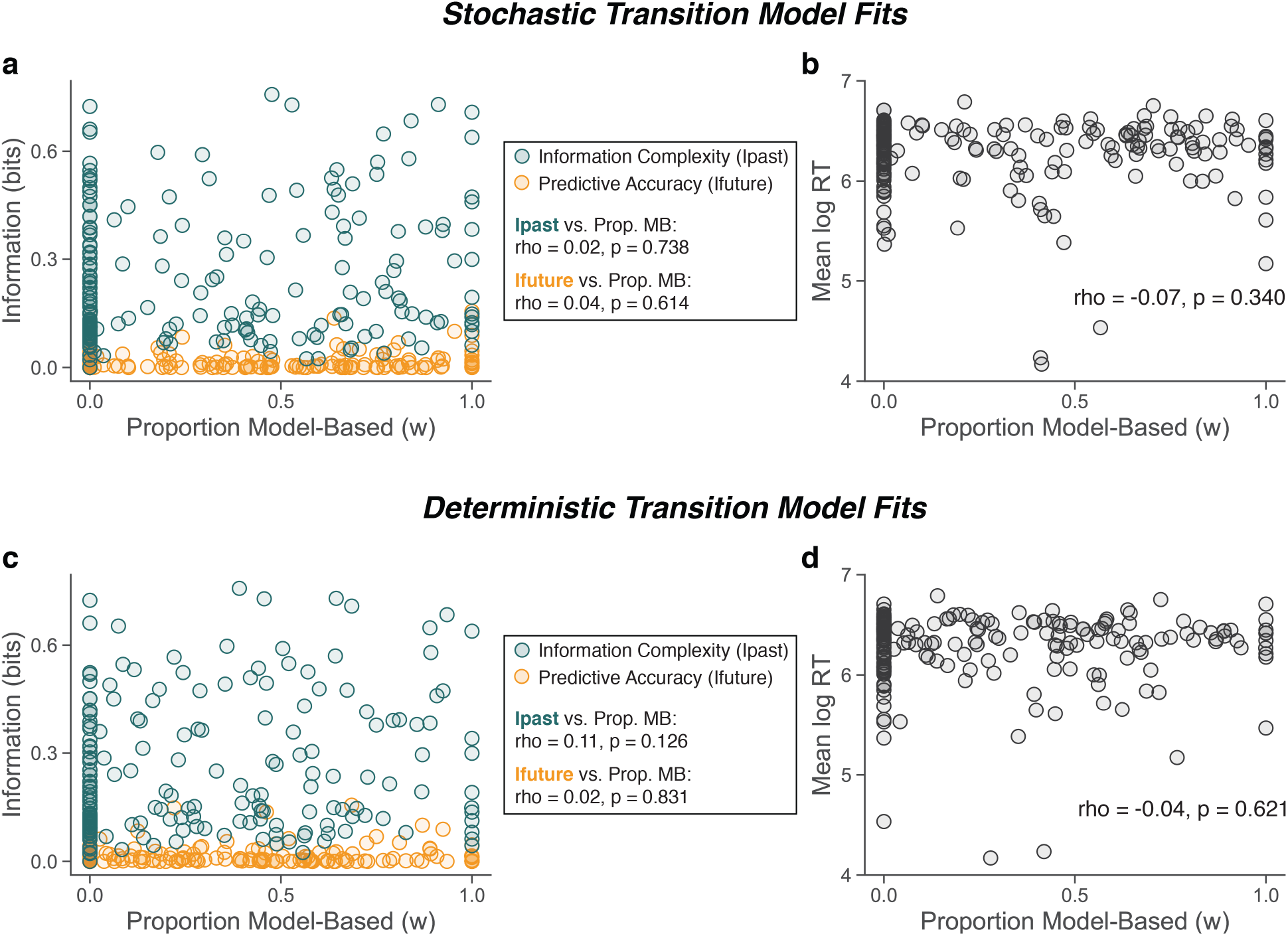
Comparison of the per-subject (circles) proportion model-based (MB) responses (best-fitting *w* parameter in the stochastic and deterministic mixture models) and: (a,c) information complexity (green) or predictive accuracy (orange), and (b,d) RTs. *rho* indicates Spearman’s correlation coefficient; *p*-values are for *H*_*0*_: *rho*=0.

Thus, although we do not know the exact algorithms human subjects used to perform the two-step task, these simulations and fits yielded two useful insights that support our behavioral findings. The first is that both model-free and model-based agents can, depending on their formulation, have relatively high information complexity. This result further dissociates computational complexity (which is higher in the model-based algorithm) from information complexity. The second is that information complexity depends critically on the specific task features each strategy uses to learn from the past and guide future choices, which for the model-free agent tended to be the observed first-step responses and for the model-based agent tended to be the latent transition probabilities between first-step responses and second-step states.

## Discussion

We measured differences in complexity between model-based and model-free learning strategies for a canonical learning task and report four main findings. First, human subject tendencies towards either model-free or model-based strategies were both associated with higher information complexity, a measure of the flexibility with which different patterns of past observations were used to guide choices. Second, while these increases in information complexity did not affect average reward obtained, they were associated with choices that had higher predictive accuracy of subsequent task features. Third, these increases in information complexity were associated with longer RTs, irrespective of whether the source of increased information complexity was based on a model-free or model-based strategy. Fourth, these similarities were apparent despite the two strategy types using very different task features to drive learning: subjects using more model-free strategies tended to learn by associating past first responses with reward, whereas subjects using more model-based strategies tended to use the inferred, latent transition probabilities between the first and second states.

Our results highlight the importance of distinguishing different forms of complexity, in particular distinguishing computational complexity, which measures resource demands, from information complexity, which measures strategic flexibility. These two forms of complexity covary under some, but not all, conditions. For example, increasing computational complexity by providing more computational resources can, in principle, support increased flexibility in how information is processed and therefore higher information complexity if these computations are used to expand the feature space over which inference is performed (Feldman & Crutchfield, 1998; Griffiths & Tenenbaum, 2003). However, as we showed, this relationship does not always hold: a standard model-free learning algorithm, although less computationally complex, can be equally or more information complex relative to a standard model-based learning algorithm.

More generally, our analyses of information complexity oppose the notion that model-free decision strategies are necessarily more automatic, inflexible, and habitual than model-based strategies, at least for human subjects performing the two-step task (Daw et al., 2011; Decker et al., 2016; Eppinger et al., 2013; Gläscher et al., 2010; Pauli et al., 2018). Both model-based and model-free strategies were associated with increases in information complexity, even if the specific information being encoded differed. Moreover, increases in information complexity for both model-free or model-based strategies were accompanied by similar increases in mean RT. Given that shorter RTs have been associated with habitual, automatic processing, such as for certain forms of statistical learning (Filipowicz, Anderson, & Danckert, 2014; Jabar, Filipowicz, & Anderson, 2017a, 2017b; Nissen & Bullemer, 1987; Robertson, 2007; Turk-Browne, Jungé, & Scholl, 2005), these results imply that the primary difference between model-free versus model-based strategies is not automaticity versus flexibility. Instead, both strategies can make flexible use of past task features to guide future behavior, but model-free strategies focus more on observed task features (e.g., responses, rewards, stimuli), whereas model-based strategies focus more on latent task features (e.g., transition structures).

Our results also highlight the usefulness of information-based metrics for assessing the nature of the strategy used by individual subjects. Although these metrics do not require an explicit model of the underlying strategy, we showed that they can be used to identify specific task features that drive learning. As predicted by a model-free algorithm, subjects with strong model-free tendencies tended to rely on their previous first-level choice. Conversely, subjects with strong model-based tendencies tended to rely on the inferred transition between the first and second states. However, those subjects also relied on the previous first-level choices to a degree that was not predicted by standard model-based algorithms. This discrepancy likely reflects a high diversity of model-based strategies used by participants for this task, which can appear similarly model-based even if they differ substantially from the common model-based algorithm used to fit their responses (Akam et al., 2015; da Silva & Hare, 2020). Moreover, there are likely substantial individual differences in the exact nature of these strategies, making it even more difficult to assess their computational complexity (da Silva & Hare, 2020). A more extensive information-based analysis of the features used by subjects on these kinds of tasks to drive learning might help inform our understanding of the specific model-based strategies they use.

Moreover, our results highlight potential future uses of the information bottleneck for assessing performance optimality across a range of strategies (Tavoni, Balasubramanian, & Gold, 2019). A strong feature of the information bottleneck method is that it can in principle compute an upper bound on the maximum achievable predictive accuracy for any given level of information complexity, and this without requiring explicit knowledge of the strategy itself (Gilad-Bachrach et al., 2003; Palmer et al., 2015; Tishby et al., 2000). This approach differs from previous approaches that assessed optimality for this task in terms of average payouts but do not take into account the nature and amount of information used by subjects to achieve those payouts (Kool et al., 2016). However, this upper bound can be difficult to compute, particularly for tasks such as the two-step task in which observations depend on subject responses. Nevertheless, future work should aim to better understand relationships between information complexity, predictive accuracy, and optimality, including how their balance is controlled by different individuals under different task conditions. New insights are likely to come from the kinds of information-bottleneck analyses that have been used previously to evaluate complexity-optimality tradeoffs in machine learning (Gilad-Bachrach et al., 2003; Tishby & Zaslavsky, 2015) and biological systems (Palmer et al., 2015). Moreover, this kind of analysis provides a strong framework in which to study notions of bounded rationality (Gigerenzer & Gaissmaier, 2011; Simon, 1955) and resource rational decision-making (Lieder & Griffiths, 2019; Tavoni, Doi, Pizzica, Balasubramanian, & Gold, 2019) that are becoming more prominent in assessing the rationality of human decision-making.

In summary, our results show that model-free and model-based learning strategies, often described as representing different ends of a continuum of information-processing flexibility, instead can be quite similar in terms of how much, how effectively, and how quickly they process information to perform a canonical learning task. These results imply that rather than distinguishing the flexibility of different learning processes, akin to the distinctions between automatic and deliberative or habitual and goal-directed processing that are often ascribed to these strategies, tasks such as the two-step task may instead distinguish between strategies that are equally complex but learn from different task features. A better understanding of these distinctions will help understand how and when these processes should be expected to vary across different healthy and psychiatric populations.

## Acknowledgements

The authors thank Wouter Kool for giving permission to use the human subject data and for providing clear, complete, and easily reproducible simulation code. We also thank Adrian Radillo for insightful comments on the manuscript and Songhan Zhang for help with simulations. Funded by NSF-NCS 1533623, R01 EB026945, and NIMH F32 MH117924. The funders had no role in the study design, data collection and analysis, decision to publish, or preparation of the manuscript.

## Author Contributions

A.L.S.F., J.L., and E.P. adapted the information bottleneck measure to the two step task; J.L. and A.L.S.F. performed the model simulations; A.L.S.F performed the model-fitting; A.L.S.F. and J.L. analyzed the human and simulation performance; G.T. and A.L.S.F. performed the algorithmic complexity analyses; all authors interpreted the results and drafted and revised the manuscript.

## Data availability

Subject data that support the findings in this study are available at: https://osf.io/z3bpk/

## Code availability

All simulations were performed without modification, using the code provided by Kool and colleagues (Kool et al., 2016; https://github.com/wkool/tradeoffs). Code for all additional subject analyses can be found at: https://osf.io/z3bpk/.

## References

Akam, T., Costa, R., & Dayan, P. (2015). Simple Plans or Sophisticated Habits? State, Transition and Learning Interactions in the Two-Step Task. PLoS Computational Biology, 11(12), 1–25. https://doi.org/10.1371/journal.pcbi.1004648

Bellman, R. (1961). Adaptive Control processes: A guided tour. Princeton, NJ, USA: Princeton University Press.

Bialek, W., Nemenman, I., & Tishby, N. (2001). Predictability, Complexity, and Learning. Neural Computation, 13(11), 2409–2463.

Bossaerts, P., & Murawski, C. (2017). Computational Complexity and Human Decision-Making. Trends in Cognitive Sciences, 21(12), 917–929. https://doi.org/10.1016/j.tics.2017.09.005

Bossaerts, P., Yadav, N., & Murawski, C. (2019). Uncertainty and computational complexity. Philosophical Transactions of the Royal Society of London. Series B, Biological Sciences, 374(1766), 20180138. https://doi.org/10.1098/rstb.2018.0138

Cormen, T., Leiserson, C., Rivest, R., & Stein, C. (2009). Introduction to algorithms. Cambridge, MA, USA: MIT PRess.

da Silva, C. F., & Hare, T. A. (2020). Humans primarily use model-based inference in the two-stage task. Nature Human Behaviour. https://doi.org/10.1038/s41562-020-0905-y

Daw, N. D., Gershman, S. J., Seymour, B., Dayan, P., & Dolan, R. J. (2011). Model-based influences on humans’ choices and striatal prediction errors. Neuron, 69(6), 1204–1215. https://doi.org/10.1016/j.neuron.2011.02.027

Daw, N. D., Niv, Y., & Dayan, P. (2005). Uncertainty-based competition between prefrontal and dorsolateral striatal systems for behavioral control. Nature Neuroscience, 8(12), 1704–1711. https://doi.org/10.1038/nn1560

Decker, J. H., Otto, A. R., Daw, N. D., & Hartley, C. A. (2016). From Creatures of Habit to Goal-Directed Learners: Tracking the Developmental Emergence of Model-Based Reinforcement Learning. Psychological Science, 27(6), 848–858. https://doi.org/10.1177/0956797616639301

Doll, B. B., Simon, D. A., & Daw, N. D. (2012). The ubiquity of model-based reinforcement learning. Current Opinion in Neurobiology, 22(6), 1075–1081. https://doi.org/10.1016/j.conb.2012.08.003

Eppinger, B., Walter, M., Heekeren, H. R., & Li, S. C. (2013). Of goals and habits: Age-related and individual differences in goal-directed decision-making. Frontiers in Neuroscience, 7(7 DEC), 1–14. https://doi.org/10.3389/fnins.2013.00253

Feldman, D. P., & Crutchfield, J. P. (1998). Measures of statistical complexity: Why? Physics Letters, Section A: General, Atomic and Solid State Physics, 238(4-5), 244–252. https://doi.org/10.1016/S0375-9601(97)00855-4

Filipowicz, A., Anderson, B., & Danckert, J. (2014). Learning what from where: Effects of spatial regularity on nonspatial sequence learning and updating. Quarterly Journal of Experimental Psychology, 67(7). https://doi.org/10.1080/17470218.2013.867518

Filipowicz, A., Anderson, B., & Danckert, J. (2016). Adapting to change: The role of the right hemisphere in mental model building and updating. Canadian Journal of Experimental Psychology/Revue Canadienne de Psychologie Expérimentale, 70(3), 201–218. https://doi.org/10.1037/cep0000078

Filipowicz, A., Glaze, C. M., Kable, J. W., & Gold, J. I. (2020). Pupil diameter encodes the idiosyncratic, cognitive complexity of belief updating. ELife, 9, e57872. https://doi.org/10.7554/eLife.57872

Gershman, S. J. (2016). Empirical priors for reinforcement learning models. Journal of Mathematical Psychology, 71, 1–6. https://doi.org/10.1016/j.jmp.2016.01.006

Gigerenzer, G., & Gaissmaier, W. (2011). Heuristic decision making. Annual Review of Psychology, 62, 451–482. https://doi.org/10.1146/annurev-psych-120709-145346

Gilad-Bachrach, R., Navot, A., & Tishby, N. (2003). An information theoretic tradeoff between complexity and accuracy. Lecture Notes in Artificial Intelligence (Subseries of Lecture Notes in Computer Science), 2777, 595–609. https://doi.org/10.1007/978-3-540-45167-9_43

Gillan, C. M., Otto, A. R., Phelps, E. A., & Daw, N. D. (2015). Model-based learning protects against forming habits. Cognitive, Affective and Behavioral Neuroscience, 15(3), 523–536. https://doi.org/10.3758/s13415-015-0347-6

Gläscher, J., Daw, N., Dayan, P., & O’Doherty, J. P. (2010). States versus rewards: Dissociable neural prediction error signals underlying model-based and model-free reinforcement learning. Neuron, 66(4), 585–595. https://doi.org/10.1016/j.neuron.2010.04.016

Glaze, C. M., Filipowicz, A., Kable, J., Balasubramanian, V., & Gold, J. (2018). A bias-variance trade-off governs individual differences in on-line learning in an unpredictable environment. Nature Human Behaviour, 2(3), 213–224. https://doi.org/10.1038/s41562-018-0297-4

Griffiths, T. L., & Tenenbaum, J. B. (2003). Probability, algorithmic complexity, and subjective randomness. Proceedings of the Annual Meeting of the Cognitive Science Society, 25(25), 480–485.

Grünwald, P., & Rissanen, J. (2007). The Minimum Description Length Principle.

Jabar, S. B., Filipowicz, A., & Anderson, B. (2017a). Knowing where is different from knowing what: Distinct response time profiles and accuracy effects for target location, orientation, and color probability. Attention, Perception, and Psychophysics, 79(8), 2338–2353. https://doi.org/10.3758/s13414-017-1412-8

Jabar, S. B., Filipowicz, A., & Anderson, B. (2017b). Tuned by experience: How orientation probability modulates early perceptual processing. Vision Research, 138. https://doi.org/10.1016/j.visres.2017.07.008

Kim, D., Park, G. Y., O’Doherty, J. P., & Lee, S. W. (2018). Task complexity interacts with state-space uncertainty in the arbitration between model-based and model-free learning. BioRxiv, 1–34. https://doi.org/10.1101/393983

Kool, W., Cushman, F. A., & Gershman, S. J. (2016). When Does Model-Based Control Pay Off? PLoS Computational Biology, 12(8), 1–34. https://doi.org/10.1371/journal.pcbi.1005090

Kool, W., Gershman, S. J., & Cushman, F. A. (2017). Cost-Benefit Arbitration Between Multiple Reinforcement-Learning Systems. Psychological Science, 28(9), 1321–1333. https://doi.org/10.1177/0956797617708288

Kool, W., Gershman, S. J., & Cushman, F. A. (2018). Planning Complexity Registers as a Cost in Metacontrol. Journal of Cognitive Neuroscience, 30(10), 1391–1404. https://doi.org/10.1162/jocn_a_01263

Lieder, F., & Griffiths, T. L. (2019). Resource-rational analysis: understanding human cognition as the optimal use of limited computational resources. Behavioral and Brain Sciences, 1–85. https://doi.org/10.1017/S0140525X1900061XP

Myung, I. J., Balasubramanian, V., & Pitt, M. A. (2000). Counting probability distributions: Differential geometry and model selection. Proceedings of the National Academy of Sciences, 97(21), 11170–11175. https://doi.org/10.1073/pnas.170283897

Nassar, M. R., Wilson, R. C., Heasly, B., & Gold, J. I. (2010). An approximately Bayesian delta-rule model explains the dynamics of belief updating in a changing environment. The Journal of Neuroscience : The Official Journal of the Society for Neuroscience, 30(37), 12366–12378. https://doi.org/10.1523/JNEUROSCI.0822-10.2010

Nissen, M. J., & Bullemer, P. (1987). Attentional Requirements of Learning : Performance Measures Evidence from. Cognitive Psychology, 19, 1–32.

O’Reilly, J. X. (2013). Making predictions in a changing world-inference, uncertainty, and learning. Frontiers in Neuroscience, 7(7 JUN), 1–10. https://doi.org/10.3389/fnins.2013.00105

Palmer, S. E., Marre, O., Berry, M. J., & Bialek, W. (2015). Predictive information in a sensory population. Proceedings of the National Academy of Sciences, 112(22), 6908–6913. https://doi.org/10.1073/pnas.1506855112

Pauli, W. M., Cockburn, J., Pool, E. R., Pérez, O. D., & O’Doherty, J. P. (2018). Computational approaches to habits in a model-free world. Current Opinion in Behavioral Sciences, 20, 104–109.

Polonio, L., Di Guida, S., & Coricelli, G. (2015). Strategic sophistication and attention in games: An eye-tracking study. Games and Economic Behavior, 94, 80–96. https://doi.org/10.1016/j.geb.2015.09.003

Robertson, E. M. (2007). The serial reaction time task: implicit motor skill learning? The Journal of Neuroscience : The Official Journal of the Society for Neuroscience, 27(38), 10073–10075. https://doi.org/10.1523/JNEUROSCI.2747-07.2007

Sebold, M., Deserno, L., Nebe, S., Schad, D. J., Garbusow, M., Hägele, C., … Huys, Q. J. M. (2014). Model-based and model-free decisions in alcohol dependence. Neuropsychobiology, 70(2), 122–131. https://doi.org/10.1159/000362840

Simon, H. A. (1955). A Behavioral Model of Rational Choice. The Quarterly Journal of Economics, 69(1), 99–118. Retrieved from http://www.jstor.org/stable/1884852

Stöttinger, E., Filipowicz, A., Danckert, J., & Anderson, B. (2014). The effects of prior learned strategies on updating an opponent’s strategy in the rock, paper, scissors game. Cognitive Science, 38(7), 1482–1492. https://doi.org/10.1111/cogs.12115

Sutton, R., & Barto, A. (1998). Introduction to reinforcement learning. Cambridge, MA: MIT Press.

Tavoni, G., Balasubramanian, V., & Gold, J. I. (2019). What is optimal in optimal inference? Current Opinion in Behavioral Sciences, 29, 117–126. https://doi.org/10.1016/j.cobeha.2019.07.008

Tavoni, G., Doi, T., Pizzica, C., Balasubramanian, V., & Gold, J. I. (2019). The complexity dividend: when sophisticated inference matters. BioRxiv, 563346. https://doi.org/10.1101/563346

Tenenbaum, J. B., Kemp, C., Griffiths, T. L., & Goodman, N. D. (2011). How to grow a mind: statistics, structure, and abstraction. Science (New York, N.Y.), 331(6022), 1279–1285. https://doi.org/10.1126/science.1192788

Tishby, N., Pereira, F. C., & Bialek, W. (2000). The information bottleneck method. ArXiv Preprint Physics/0004057, 1–16. https://doi.org/10.1108/eb040537

Tishby, N., & Zaslavsky, N. (2015). Deep learning and the information bottleneck principle. 2015 IEEE Information Theory Workshop, ITW 2015, 1–5. https://doi.org/10.1109/ITW.2015.7133169

Turk-Browne, N. B., Jungé, J., & Scholl, B. J. (2005). The automaticity of visual statistical learning. Journal of Experimental Psychology. General, 134(4), 552–564. https://doi.org/10.1037/0096-3445.134.4.552

Voon, V., Derbyshire, K., Rück, C., Irvine, M. A., Worbe, Y., Enander, J., … Bullmore, E. T. (2015). Disorders of compulsivity: A common bias towards learning habits. Molecular Psychiatry, 20(3), 345–352. https://doi.org/10.1038/mp.2014.44

